# Confocal reflectance microscopy for mapping collagen fiber organization in the vitreous gel of the eye

**DOI:** 10.1101/2022.11.09.515466

**Authors:** Eileen S. Hwang, Denise J. Morgan, Jieliyue Sun, M. Elizabeth Hartnett, Kimani C. Toussaint, Brittany Coats

## Abstract

Vitreous collagen structure plays an important role in ocular mechanics, but capturing this structure with existing vitreous imaging methods is hindered by loss of sample position and orientation, low resolution, or small field of view. The objective of this study was to evaluate confocal reflectance microscopy as a solution to these limitations. The use of intrinsic reflectance avoids staining and optical sectioning eliminates the requirement for thin sectioning, minimizing processing for optimal preservation of natural structure. We developed a sample preparation and imaging strategy using *ex vivo* grossly sectioned porcine eyes. Imaging revealed a network of uniform diameter crossing fibers (1.1 ± 0.3 μm for a typical image) with generally poor alignment (alignment coefficient = 0.40 ± 0.21 for a typical image). To test the utility of our approach for detecting differences in fiber spatial distribution, we imaged eyes every 1 mm along an anterior-posterior axis originating at the limbus and quantified the number of fibers in each image. Fiber density was higher anteriorly near the vitreous base, regardless of imaging plane. These data demonstrate that confocal reflectance microscopy addresses the previously unmet need for a robust, micron-scale technique to map features of collagen networks *in situ* across the vitreous.

## 1. Introduction

The vitreous, a transparent gel between the lens and retina of the eye, plays a fundamental role in retinal diseases such as retinal detachment and macular hole. In these diseases, the vitreous exerts traction on the retina. Given the importance of collagen fiber organization and structure on the mechanics of many biomaterials [1], the ability of the vitreous to transmit forces to the retina, generated by eye movements and gravity, likely also depends on the structure of the collagen fibers within the vitreous. To evaluate the effect of collagen structure on traction force distribution, Aravelo et al. created artificial collagen gel networks attached to microsphere-embedded polyacrylamide gels [2]. A shear force was applied to the free end of the collagen network and the resulting stress, or force, distribution was calculated on the attached polyacrylamide gel. Force distribution was highly heterogeneous and locations of higher stress were due to alignment of collagen fibers in the direction of principal loading. These data suggest that micron-scale imaging of the vitreous may provide critical clues to the force distribution on the retina and inform diagnostic and treatment strategies for traction-based retinal diseases.

To date, research into vitreous structure has been limited by the lack of an imaging technique that can depict microscopic structure as well as higher-level organization and spatial trends. To section the vitreous, it must be made rigid, but processing the vitreous to make it rigid is highly problematic since it is 99.8% water. Embedding requires dehydration and results in artifactual shrinkage and loss of the central vitreous [3]. Freezing extracellular matrix gels creates a honeycomb-like artifact [4]. Although plastic embedding and rapid freezing have been used to minimize these effects during vitreous imaging [5,6], artifacts may not be completely eliminated.

Two techniques have been used to image vitreous structures without processing: slit-lamp biomicroscopy and phase contrast microscopy. Slit-lamp biomicroscopy captures an optical cross-section of the vitreous body by imaging side-scattered light against a dark background. Slit-lamp images show alternate bands of high scattering and low scattering in human vitreous, but this technique results in low resolution images that lack information about the microscopic structure needed to understand the origin of the mechanical properties of the vitreous [7]. The other processing-free technique, phase contrast microscopy, relies upon constructive interference to provide contrast in unstained samples. This method provides high resolution images of ~1 μm fibers in the vitreous [8]. However, phase contrast microscopy is a transmission-based technique without optical sectioning, and thus requires cutting out a small piece of the soft vitreous gel and allowing it to deform into a flat layer, thereby losing information about the original shape, location, and orientation.

Confocal microscopy is a promising approach for viewing the natural structural organization of the vitreous because it is a high resolution method that relies on optical sectioning rather than physical sectioning. A specific type of confocal microscopy, confocal reflectance microscopy, can be used to image samples with minimal processing since it does not require staining. Unlike confocal fluorescence, which detects light at a wavelength longer than that used to illuminate the sample, confocal reflectance relies upon detection of light at the same wavelength as the laser used to illuminate the sample. Confocal reflectance microscopy of artificial collagen gels shows that the density and orientation of the micron-scale structures imaged by this technique are highly correlated with material properties such as gel stiffness [1]. This suggests that vitreous structures imaged by this technique will provide insight into vitreous material properties. Confocal reflectance microscopy has been used to image cells in the cornea in live humans [10], but has yet to be applied to the vitreous to the best of our knowledge.

In summary, many techniques have been applied to investigate vitreous structure, but there remains an unmet need for a fiber imaging method that preserves natural fiber organization in the vitreous and maps images to specific locations within the eye. Quantitative mapping of fiber network features across the eye is crucial to inform computational, conceptual, and physical models of the vitreous in health and disease. In this work, we evaluate whether confocal reflectance microscopy can fulfill the need for mapping micron-scale collagen fiber organization in the vitreous in its native state.

First, we develop and describe methods for sample preparation, image acquisition, and image analysis. These methods enable us to provide the first micron-scale images of the collagen fiber network of the vitreous *in situ* without staining or processing. We then vary key parameters in sample preparation, such as delay times and sample treatments, to determine the limits that must be respected to minimize artifacts. Next, we apply spatial mapping to answer the open question of whether fiber density, width, and orientation vary in specific regions, planes, and axes within the eye. Finally, we examine how our findings correlate with findings from previously described methods for evaluating spatial differences in vitreous structure.

## 2. Methods

Our initial set of experiments sought to qualitatively evaluate the robustness of confocal reflectance imaging of the vitreous to various alterations in sample processing. Once the optimal parameters for sample preparation were determined, we applied automated fiber segmentation to extract information about density, width, and orientation. We compared the automated fiber segmentation method to alternate methods for evaluating density and orientation. We then applied mixed effects models to assess regional and orientational trends in density, width, alignment, and angular mean.

### 2.1 Sample preparation

Studies were performed on *ex vivo* eyes from 2-4 year old female domestic pigs sourced from a meat processing plant. Porcine eyes were selected because they are similar in size to human eyes, and because they contain concentrations of collagen and hyaluronic acid on similar orders of magnitude to those in human eyes [9,10]. Eyes were only obtained from female pigs because male pigs were not available from the meat processing plant. Each pig’s right eye was selected to avoid including two eyes from the same pig. Eyes with weights greater than or less than 1 standard deviation from the mean weight were excluded.

Each eye was embedded in an agarose cube, and then sectioned in a single plane to obtain a cap (Fig. 1). The cap was placed on a large coverslip and the edges were sealed with silicone caulk to prevent the vitreous from spreading out past the sclera edge. The agarose was trimmed slightly prior to coverslip application to enable the sclera to be flush against the coverslip, minimizing trapped air.

**Fig. 1.**
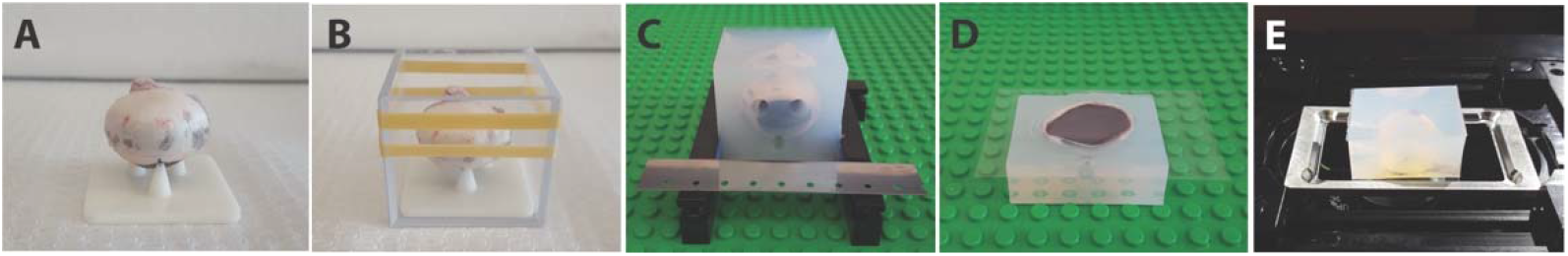
Steps of sample preparation. A, placing the eye on a three-pronged pedestal with the 6 o’clock inferior limbus on one prong. B, placing the eye in a transparent cube mold for addition of agarose. C, sectioning the eye using a microtome blade. D, application of silicone caulk and a large coverslip to the cut face of the sample. E, inverting the sample and placing it on an aluminum coverslip holder for imaging on an inverted confocal microscope.

### 2.2 Image acquisition

Confocal reflectance microscopy was performed on an Olympus FV1000 inverted microscope with a 30× 1.05 NA silicone oil objective. The imaging plane was oriented parallel to the sectioned plane to permit scanning across the ~2 cm open face of the sample with the motorized stage. Z-stacks demonstrated motion artifacts consistent with drift within the sample due to fluidity during the 3.4 s image acquisition time. Our primary objective was to evaluate differences in network properties between different locations across the eye, so we chose to eliminate this artifact by obtaining two-dimensional 70-μm × 70-μm images at a fixed depth (350 μm). At shallower depths, background noise increased and at greater depths, signal strength decreased. The sample was illuminated by a tightly focused 473-nm wavelength laser beam, and the reflected light was captured through a pinhole by the detector. The use of a pinhole segregates light coming from the plane in focus from all other out-of-focus regions. To reduce the artifacts from coverslip reflection, we minimized the confocal aperture (50 μm) and maximized laser power (15 mW). To avoid the bright center spot, we limited image acquisition to a decentered subset of the optical imaging field (Supplement 1, Fig. S1). We obtained images with a 12.5 μs dwell time and a pixel resolution of 256×256. We used Kalman averaging of three scans per line within the Olympus Fluoview software to enhance signal relative to background noise.

### 2.3 Evaluating sample storage and treatment

We qualitatively assessed the effect of postmortem time, sample freezing, sample fixation, post-dissection time, and exposure to air on the appearance of the vitreous collagen fibers as imaged by confocal reflectance microscopy. All samples were compared by sectioning the globe tangential to the inferior limbus and imaging the sample every 1 mm along the anterior-posterior axis originating at the inferior limbus. To assess the effects of postmortem time, we compared unfixed eyes imaged at 4 h postmortem, 48 h postmortem, and 1 week postmortem (2 eyes at each time point). To assess the effects of freezing, we prepared 2 eyes by immersing the entire globe in 2-methylbutane chilled in a dry ice-isopropanol bath and then sectioning the eye with a band saw. The frozen cap was placed faced down on a coverslip and allowed to thaw for 1 h prior to imaging. These eyes were compared to 2 fresh eyes at 1 h post-dissection. To assess the effect of tissue fixation, we prepared 3 fixed samples by immersing the entire intact eye in 4% paraformaldehyde for 4 h at room temperature prior to embedding in agar and sectioning. Three fresh/unfixed control samples were prepared as described above while avoiding trapped air. In addition, 3 fresh/unfixed samples were prepared with a small air bubble trapped with the vitreous in the cap. The samples were imaged periodically until 20 h after the cap was prepared. All subsequent imaging was performed on unfixed/fresh eyes within 1 h post-dissection within 48 h postmortem.

### 2.4 Collagenase digestion

To verify that the fibers imaged in the vitreous by confocal reflectance were composed primarily of collagen fibers, we tested the effect of treatment with collagenase. A 50 μL piece of vitreous was dissected from a fresh porcine eye and cut in half. Each 25 μL piece of vitreous was placed in an 8 mm diameter coverslip-bottom chamber with 200 μL of either 2 mg/mL bacterial collagenase type II (Worthington) or phosphate-buffered saline. Specimens were incubated at 37° C for 2 h on a shaker and then imaged by scanning the entire sample chamber.

### 2.5 Quantitative image analysis

The images were first preprocessed by smoothing using the non-local means denoising Fiji plugin (settings: sigma = 20, smoothing factor = 1) [11]. Automated fiber segmentation was then performed with the Fiji ridge detection plugin (settings: line width = 5, high contrast = 115, low contrast = 60, method for overlap resolution = slope, minimum line length = 10) [12]. The optimal value for each parameter was determined by varying the parameter on a test image followed by visual review of the fiber segments drawn on the image by the plugin preview function. The tabular output from the ridge detection consisted of variable length fiber segments which were stitched together using a custom MATLAB script developed by Creveling et al. [13]. This script removes duplicate fibers and determines fiber orientation and width. Fiber density was defined as the total number of unique fibers in each −70 μm × 70-μm image.

Fiber orientation was determined by fitting a line through the segment endpoints and quantifying the acute angle of that line relative to the horizontal axis of the image. Given that a fiber does not have an origin, i.e. the beginning and end are indistinguishable, the orientation was measured on a scale from −90 degrees to +90 degrees. For each image, the distribution of orientations was used to determine the alignment coefficient, which ranged from 0 (not aligned) to 1 (perfectly aligned). The alignment coefficient was defined as the magnitude of the mean of the unit vectors created by the fiber, which was calculated using the CircStat Matlab package [14]. The width of a single fiber was defined by averaging the segment widths along the length of the fiber.

To test the ability of our custom segmentation method to adequately quantify collagen fiber density, two alternate methods were used to quantify fiber density in the subset of images obtained from the axis originating at the nasal limbus imaged in the tangential plane: manual fiber counting and automated pixel thresholding (Fig. 2). Manual fiber counting was performed by printing the image at a set size and tracing each fiber to count the number of fibers per image. At fiber intersections, the fibers were assumed to continue their original orientation. Preliminary experiments demonstrated high correlation of manual fiber counting between two masked graders, however the final analysis was performed by only one grader. Automated pixel thresholding was performed by turning the greyscale image into a binary image using a threshold automatically determined by the Fiji moments algorithm. The number of white pixels per image was summed to provide a percentage of pixels occupied by signal.

**Fig. 2.**
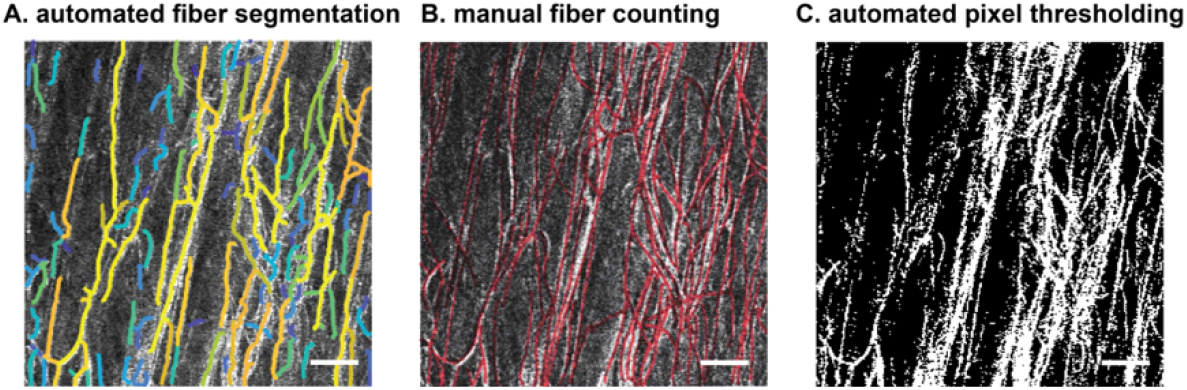
Illustration of three different methods for determining fiber density in an example image. Scale bar = 10 μm.

To test the ability of our custom segmentation method to quantify fiber alignment, the Fiji OrientationJ plugin was used as an alternate method to calculate orientation and alignment. A structure tensor containing directional information is calculated by the weighted inner product of partial spatial derivatives of the image. The local orientation and isotropy properties (gradient energy and coherency) can be easily derived from this structure tensor. A structure tensor, a matrix representative of the partial derivatives of the image, is calculated using a Gaussian weighting function. The structure tensor is then used to calculate the orientation, energy, and coherency for each pixel. Coherency is defined as the ratio between the difference and sum of the tensor eigenvalues. Similar to the alignment coefficient, coherency represents a circular spread in fiber orientations for each image and varies from 0 (not aligned) to 1 (perfectly aligned). The dominant fiber direction was also extracted from the histogram of orientations in OrientationJ to compare to the mean angle from our custom fiber segmentation method.

### 2.6 Spatial variation of fiber density, width and alignment

To determine whether confocal reflectance could be used to detect spatial variation in fiber features, we tested whether images obtained from near the vitreous base (in the anterior portion of the eye by the limbus) differed from images obtained more posteriorly. This was first evaluated this question by imaging 5 eyes sectioned in a plane tangential to the nasal corneal limbus at 3 o’clock every 1 mm along the anterior-posterior axis originating at the nasal limbus (Fig. 3A, B). The imaging plane was parallel to the sectioning plane at a constant depth of 350 μm. Images without fibers on visual inspection were excluded, and eyes with greater than 2 excluded images were eliminated from the analysis. Next, to evaluate the effects of the imaging plane, we imaged each location along the anterior-posterior axis originating at the nasal limbus in 4 eyes sectioned in a radial plane (Fig. 3C,D).

**Fig. 3.**
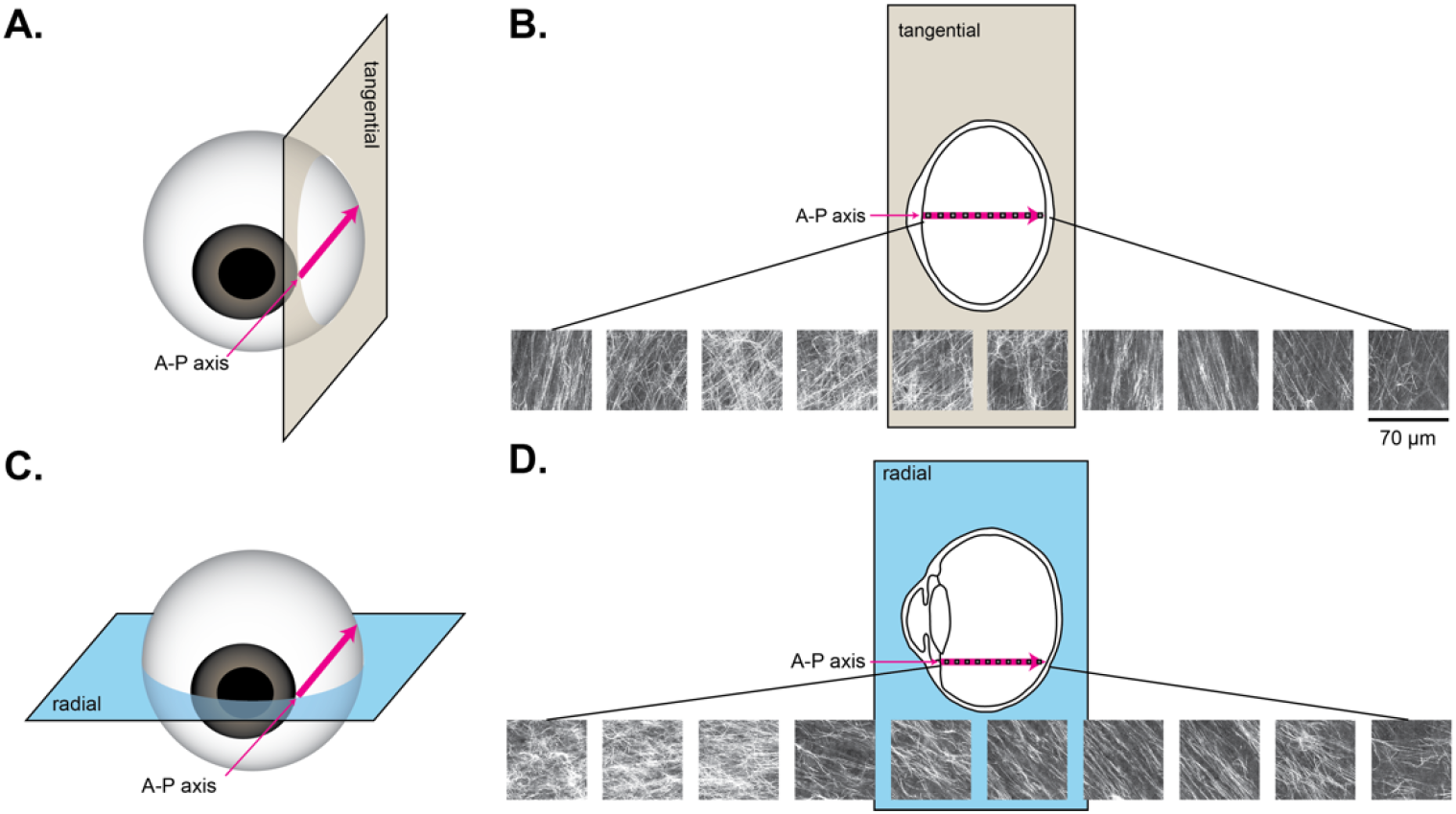
Illustration of two orthogonal planes intersecting at the anterior-posterior (A-P) axis originating at the nasal (3 o’clock) limbus on the right porcine eye. Imaging was performed in both planes at equivalent locations every 1 mm along the A-P axis at a constant depth (350 μm). A, Schematic illustrating the imaging plane tangential to the corneal limbus. B, Example images obtained in the tangential plane. C, Schematic illustrating the imaging plane radial to the corneal limbus. D, Example images obtained in the radial plane.

Our next goal was to evaluate possible radial symmetry in collagen fiber features as suggested by low-resolution slit-lamp imaging [15]. To this end, we imaged collagen fibers in planes tangential to the limbus along anterior-posterior axes originating at the superior (12 o’clock, 7 eyes), inferior (6 o’clock, 4 eyes) and temporal limbus (9 o’clock, 5 eyes) (Fig. 4).

**Fig. 4.**
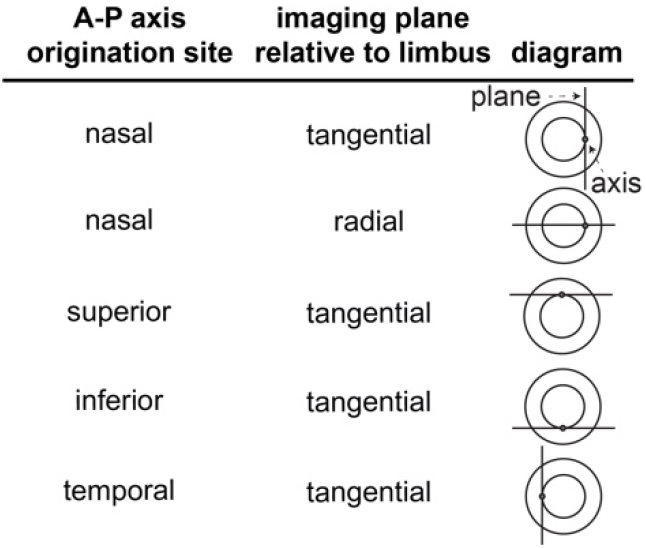
Illustration of five imaging datasets obtained along anterior-posterior (A-P) axes originating at four locations along the limbus in two imaging planes.

### 2.7 Statistical Analysis

We performed statistical analyses of three types. 1) We evaluated the association of fiber density, width, and alignment with the distance along the anterior-posterior axis. 2) We compared fiber density, width, and alignment between different imaging axis origination sites and planes. 3) We correlated fiber density, alignment, and orientation quantified by different methods.

To evaluate the association between fiber density, width, and the alignment coefficient with the distance from the inner cornea, we used a mixed effects linear regression model which accounts for clustering due to multiple measurements taken per sample. In a clustered data set, a correlation coefficient and a multiple R statistic are not defined because they are a mixture of intra-cluster correlations and inter-cluster correlations. Therefore, we report slopes rather than correlation coefficients or coefficients of determination. In each case, we evaluated the association between the outcome variable and the distance from the inner cornea in the 5 datasets: superior, nasal-tangential, nasal-radial, inferior, and temporal (Fig. 4). To account for multiplicity in evaluating the trend in the 5 datasets, we used the Hommel procedure to calculate adjusted p-values [16]. An adjusted p-value <0.05 was considered significant. All statistical evaluations were performed using STATA version 16.1 or GraphPad Prism 9.4.1.

Next, we evaluated whether fiber density, width, and alignment coefficient differed between the 5 datasets using multiple pairwise comparisons with adjusted p-values. To obtain means and standard errors that correctly account for clustering within data, we fit a mixed effects linear regression model with the image dataset and the distance from the inner cornea as the two predictor variables and then obtained the means and standard errors using post-fit marginal estimation.

Finally, we evaluated the correlation between different methods of measuring fiber density, fiber alignment, and fiber orientation. We chose data from a single sample to calculate the correlation coefficient since there is no cluster effect if there is only one cluster in the data analyzed. We used a Pearson correlation and defined strong correlation as R > 0.7.

## 3. Results

### 3.1 Imaging results

The confocal reflectance images of the porcine vitreous revealed a network of crossing fibers (Fig. 5A).

**Fig. 5.**
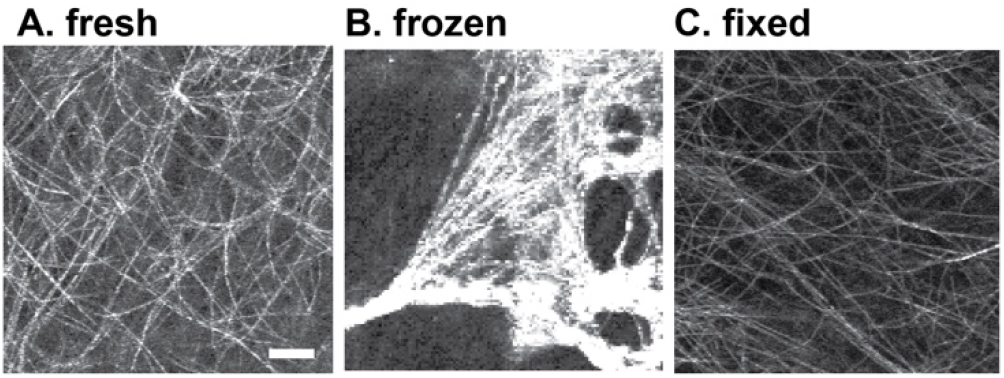
Confocal reflectance images of fresh vitreous (A), frozen/thawed vitreous (B), and fixed vitreous (C). Scale bar = 10 μm.

### 3.2 Sample storage and treatment

To determine the optimal method of sample preparation, we evaluated the effects of increased postmortem time, freezing, and fixation on the appearance of the vitreous by confocal reflectance. There was no observable difference in vitreous collagen fibers in eyes where dissection began at 4 h, 48 h, and 1 week postmortem (data not shown). At 1 week postmortem, retina debris was present at the interface between the vitreous and the coverslip following dissection. Freezing, sectioning and thawing samples resulted in large empty vacuoles separated by beams of thick, reflective fibers that were not present in fresh samples (Fig. 5B). Fixation of samples in 4% paraformaldehyde did not alter the appearance of the collagen fibers (Fig. 5C) compared to the fresh samples.

Given the well-established effects of post-dissection time and exposure to air on the mechanical properties of the vitreous, we hypothesized that structural changes can be detected using confocal reflectance after dissection and exposure to air. Specifically, we tested whether the vitreous appearance was altered by incubating the sample after preparing the cap on the coverslip. We also tested whether air trapped within the vitreous cap accelerated these alterations, and whether fixation inhibited these alterations. We found that increased time between dissection and imaging resulted in fibers of greater reflectance intensity and width (Fig. 6).

**Fig. 6.**
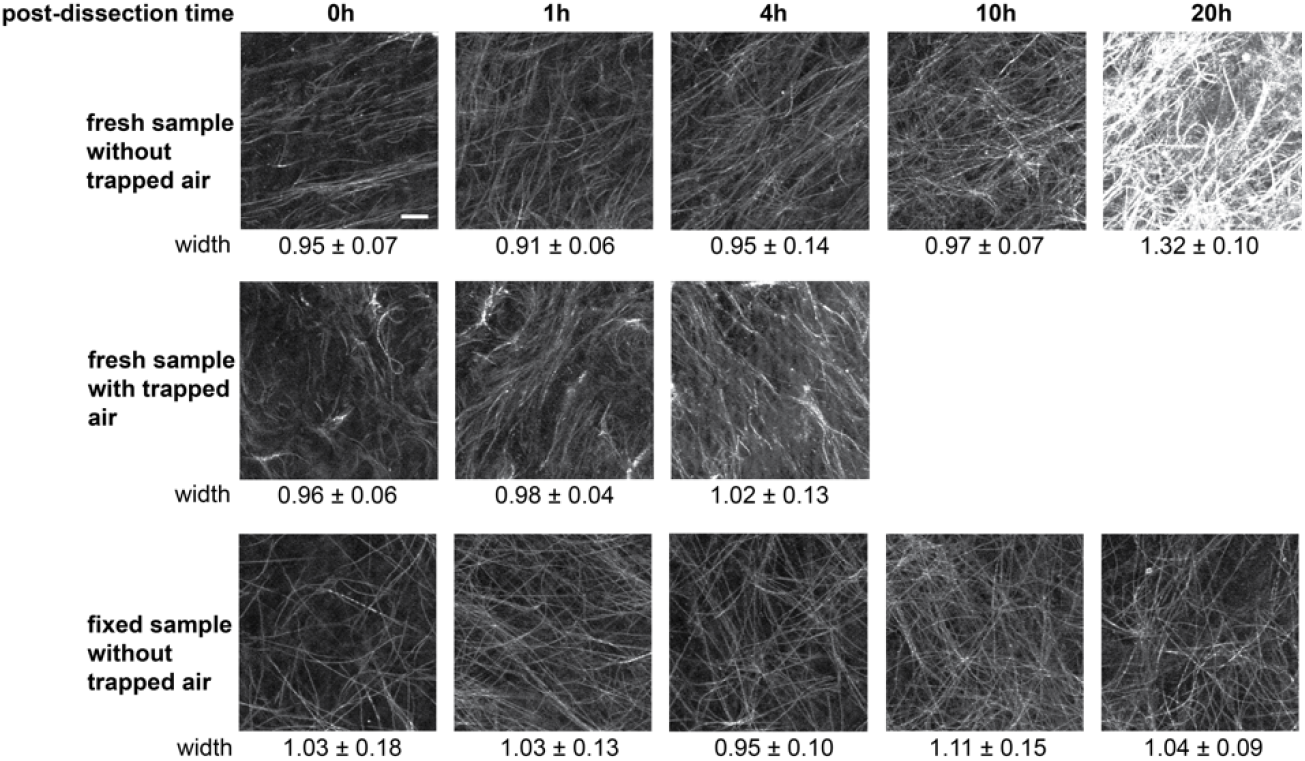
Changes in vitreous fiber appearance over time post-dissection in fresh samples without a trapped air bubble (top row), fresh samples with an air bubble trapped during cover slip application (middle row), and fixed samples without a trapped air bubble (bottom row). Width is mean ± SD from 3 replicates. Scale bar = 10 μm.

These changes were minimal during the first 4 h in the samples without trapped air, suggesting that imaging within 4 h minimizes post-dissection effects in these samples (Fig. 6). However, when an air bubble was trapped during coverslip application, changes in fiber intensity and width occurred sooner than 4 h. Fixation using 4% paraformaldehyde prevented changes in reflectance intensity and width for at least 20 h after dissection.

To confirm that the fibers imaged were composed of collagen, we treated the vitreous with bacterial collagenase enzyme, which breaks down collagen. Fibers were visibly absent in specimens incubated with collagenase for 2 h compared to controls incubated with phosphate-buffered saline (Fig. 7).

**Fig. 7.**
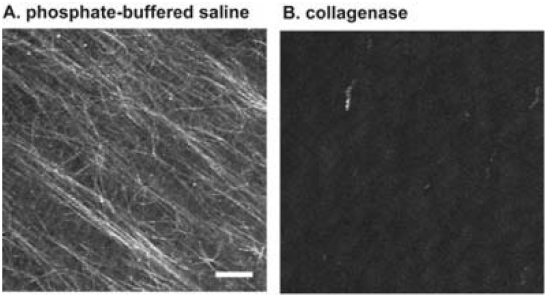
Confocal reflectance images of vitreous (A) after incubation in phosphate-buffered saline for 2 h at 37°C, and (B) in 2mg/ml collagenase for 2 h at 37°C. Scale bar = 10 μm

### 3.3 Assessment of spatial variation in fiber density

Based on previous reports of higher collagen density near the vitreous base as measured by hydroxyproline concentration after acid hydrolysis [17], we postulated that confocal reflectance microscopy would demonstrate a higher density of collagen fibers near the vitreous base. We first tested this hypothesis by imaging points at various distances from the inner cornea along an anterior-posterior axis originating at the nasal limbus in an eye sectioned tangentially to the 3 o’clock limbus (Fig. 3). Using automated fiber segmentation, a statistically significant decrease in fiber density from the front to the back of the eye was found (Fig. 8A).

**Fig. 8.**
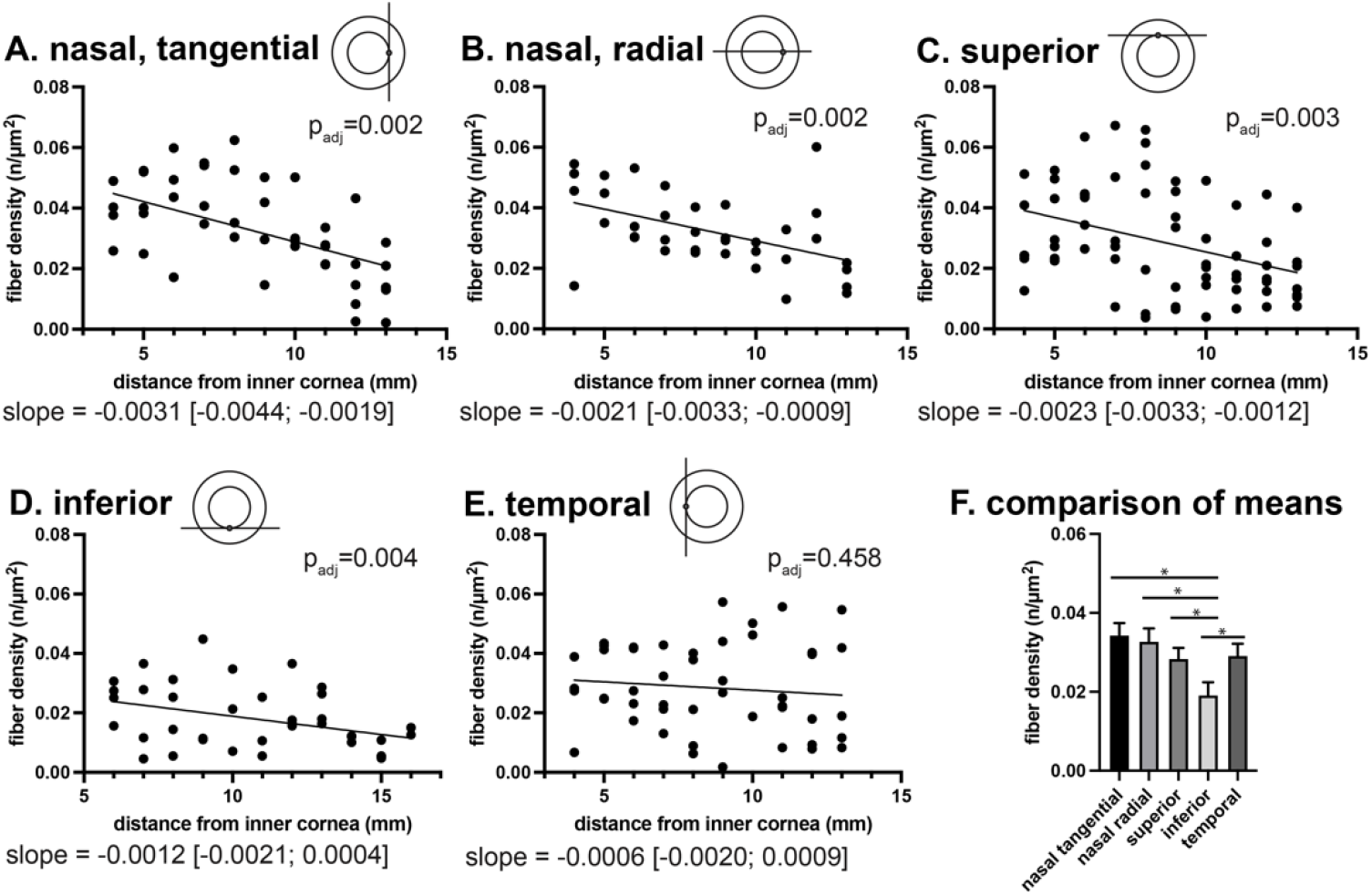
Trend in fiber density as measured by automated fiber segmentation along anterior-posterior axes originating at four different sites (A, B) nasal, (C) superior, (D) inferior, and (E) temporal, imaged in tangential planes (A, C, D, E) or a radial plane (B). Distance from the inner cornea is equivalent to distance along the A-P axis. F, comparison of overall density between the 5 datasets using a mixed effects model accounting for clustering and distance from the inner cornea. Brackets indicate 95% confidence intervals. Error bars represent standard error. Asterisks indicate p_adj_ ≤ 0.001.

To evaluate the robustness of our findings, we applied two alternate methods for assessing fiber density (Fig. 2, Supplement 1, Fig. S2). All three quantification methods found statistically significant trends along the anterior-posterior axis originating at the nasal limbus imaged in the tangential plane. Based on these results, we chose to proceed with automated fiber segmentation for additional quantitative analyses of fiber network features.

Because confocal reflectance is unable to detect fibers that are near-perpendicular to the imaging plane, we imaged locations along the anterior-posterior axis in two planes originating at the nasal limbus (radial and tangential, Fig. 3). We found a significant trend of decreasing fiber density with increasing distance from the inner cornea in both orthogonal planes (Fig. 8A, B). Fiber density (mean ± SE) in all images from the nasal-tangential dataset was 0.034 ± 0.002 fibers/μm^2^. The fiber density in all images from the nasal-radial dataset was 0.032 ± 0.002 fibers/μm^2^, which was not significantly different (p = 0.57, Fig. 8F).

Next, we hypothesized that due to radial symmetry of the eye, there would be similar density trends along anterior-posterior axes originating at the superior (12 o’clock), inferior (6 o’clock) and temporal (9 o’clock) limbus. Since density findings originating at the nasal limbus were similar when imaged in two orthogonal planes (tangential and radial), we evaluated the axes originating at other sites in a tangential plane only. We found a significant trend of decreasing density along the anterior-posterior axes originating at the superior and inferior limbus, but not at the temporal limbus (Fig. 8C, D, E). To further investigate the concept of radial symmetry, we performed pairwise comparisons of collagen fiber density from the superior, nasal-tangential, nasal-radial, inferior, and temporal datasets. We found that the inferior dataset had significantly lower density compared to the other four datasets (Fig. 8F, p_adj_≤0.001).

### 3.4 Analysis of fiber width

We found that fiber width was 1.1 ± 0.3 μm (mean ± SD) in a typical image (Fig. 5A). No sizeable (>10%) differences in width were observed with increasing distance from the inner cornea or between origination sites (Supplement 1, Fig. S3). The confocal reflectance approach allows us to determine for the first time that fiber width is relatively constant in the imaged areas of the porcine eye.

### 3.5 Analysis of fiber alignment and orientation

Based on slit-lamp images from human eyes, [15] we hypothesized that fibers would be aligned parallel to the anterior-posterior axis, regardless of the origination site. We also anticipated that overall alignment would be weak based on low retardation measured by quantitative polarized light microscopy [18]. After automated fiber segmentation, we used linear regression to obtain an angle for each fiber. To characterize the alignment between fibers in a given image, the distribution of orientations was used to calculate an alignment coefficient ranging from 0 (not aligned) to 1 (perfectly aligned). Using these methods, a wide distribution of orientations was observed in a typical image (alignment coefficient = 0.40 ± 0.21, Fig. 5A). No sizeable (>10%) trends with increasing distance from the inner cornea were noted (Supplement 1, Fig. S5). The alignment coefficient was significantly higher in the nasal-radial dataset compared to the other 4 datasets (p_adj_ < 0.0001, Fig. 9A).

**Fig. 9.**
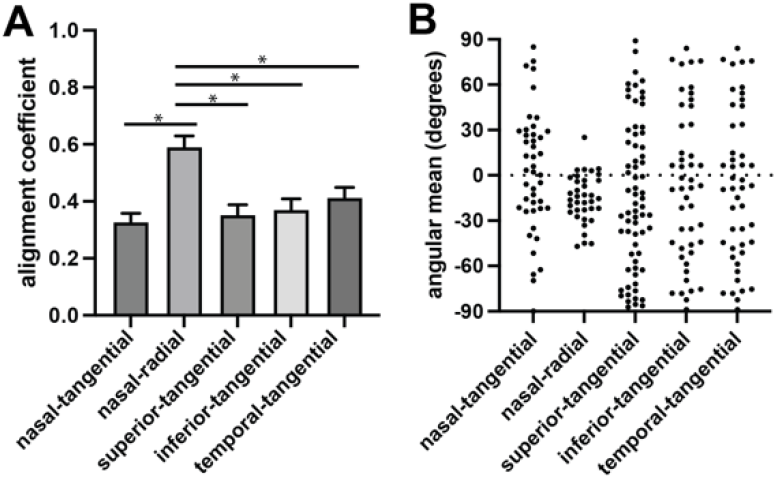
A, Comparison of alignment coefficient between the 5 datasets using a mixed effects model accounting for clustering and distance from the inner cornea. Error bars represent standard error. Asterisks indicate p_adj_ ≤ 0.0001. B, Comparison of angular means between the 5 datasets.

Based on the generally anterior-posterior orientations that Filas et al. observed by quantitative polarized light microscopy [18] along the anterior-posterior axes originating at the nasal and temporal limbus, we anticipated that we would find angular means around 0 degrees, which we defined as parallel to the anterior-posterior axis. Although the angular means in the nasal-radial dataset were closer to zero, the angular means in the remaining 4 tangential datasets were widely distributed (Fig. 9B). We confirmed the differences in orientation between nasal-radial and nasal-tangential datasets with an alternate method for analysis (Supplement 1, Fig. S7). Using a radial plane instead of a tangential plane to image the axis originating at the nasal limbus impacted orientation measurements (alignment coefficient and angular mean), although density measures were not affected (Fig. 8F).

## 4. Discussion

We demonstrated that confocal reflectance microscopy is a robust and reliable approach for mapping collagen fiber network characteristics across the vitreous gel of the eye. Using confocal reflectance, we identified a network of fibers that was robust to fixation with PFA and postmortem storage up to 48 h. In contrast, freezing created vacuoles within the vitreous, and post-dissection storage of fresh samples resulted in wider, more intensely reflective fibers, a change that occurred more quickly with exposure to air. Previous researchers found that vitreous gel stiffness changes significantly post-dissection, especially if it is exposed to air, due to exudation of a hyaluronic-acid rich fluid [19]. The correlation of post-dissection changes in material properties with post-dissection changes in fiber appearance, and prior work in artificial collagen gels correlating confocal reflectance images with material properties [1,2] suggest that vitreous fiber network properties identified by confocal reflectance are highly relevant to vitreous gel material properties [3,6,17].

Some prior vitreous imaging methods required extracting a small sample of the vitreous gel, which resulted in loss of original position, orientation, and shape of the imaged sample. In contrast, we adopted the strategy of Filas et al. of grossly sectioning the eye a single time and imaging near the surface of the open face [18,20]. This preparation strategy, combined with the large range of motion of the microscope stage, allowed us to map the images to a specific location within the eye and evaluate whether our technique was able to detect previously-established regional differences in collagen density [3,6,17]. We found the expected decrease in density along an anterior-posterior axis originating at the superior, nasal, and inferior limbus. We did not find a similar trend at the temporal limbus, which may be due to the temporally peaked shape of porcine cornea. The ability to spatially register our imaging locations allowed us to quantify trends in fiber properties along specific axes in the eye, an accomplishment that was not possible with prior techniques employed for vitreous imaging.

Confocal reflectance does not require staining or thin sectioning, and therefore avoids processing artifacts from dehydration and freezing. Stain-free imaging of vitreous collagen fibers is challenging because they scatter very little light. Past researchers were able to use phase contrast to increase contrast and image vitreous fibers after removing a small sample from the eye [21]. They found a multi-directional network of ~1-μm diameter fibers, similar to what we observe by confocal reflectance. We theorize that the confocal system is necessary to powerfully reduce speckle noise due to inherently weak signal of vitreous fibers. The inherently weak signal likely also explains why we were unable to image vitreous with second harmonic generation microscopy, even when using an ultra-sensitive, refractive-index-matched substage detector [22], and why Filas et al. were not able to image individual fibers with quantitative polarized light microscopy [18]. They were able to extract measures of average alignment and orientation, and found poor alignment, consistent with our findings. Although reflectance is beneficial because it provides contrast without staining, its major weakness is lack of reflectance from fibers that are near perpendicular to the imaging plane [23]. We addressed this weakness by imaging equivalent locations in two orthogonal planes along an anterior-posterior axis originating at the nasal limbus. We found that density was similar between the two planes, but the fibers imaged in the radial plane were significantly more aligned along the anterior-posterior axis than fibers imaged in the tangential plane. Thus, the predominance of anterior-posterior streaks seen in human vitreous by slit-lamp, and the average anterior-posterior orientation mapped along the nasal and temporal axes by quantitative polarized light microscopy may be explained by fibers oriented in the tangential plane [7,18,24].

One limitation is that our approach requires sectioning the globe. Therefore, numerous *ex vivo* eyes are needed to comprehensively image the eye along a series of representative planes. Another limitation of imaging a soft gel is sample movement, which made it impossible for us to follow fibers from layer to layer within a z-stack. Another potential limitation is that there is likely a lower detection limit for fibers. Specifically, fibers of a narrow diameter or that are composed of fibrils with weaker alignment within the fiber may not be detected by confocal reflectance. We are unable to completely exclude this as a confounder, but the correlation of our observed trends with other methods in the literature that are independent of fiber size, such as hydroxyproline content, suggests that the trend we observed reflected a trend in collagen content rather than simply fiber visibility.

## 4. Conclusion

In summary, we established an approach for applying confocal reflectance microscopy to the vitreous gel of the eye. Confocal reflectance overcomes several weaknesses of prior vitreous imaging strategies by capturing micron-scale network features with spatial registration across the entire eye and minimal processing artifact. We illustrate the validity of this technique in comparison with existing methods for assessing regional differences in collagen fiber density and orientation, probe the limitations of using reflectance, and identify boundaries of sample preparation parameters. Considering the established correlation of collagen gel microstructure with material properties, our data suggest that confocal reflectance can be used to uncover the structural basis for alterations in vitreous gel stiffness that lead to vitreoretinal interface stress, retinal detachment, and vision loss.

## Acknowledgments

We are grateful to Gregory J. Stoddard for statistical support, Dustin J. Layton for design and manufacturing of sample preparation devices, Izabela Bulska for assistance with graphic design of figures, and Alexander Dvornikov and Enrico Gratton for the opportunity to use the DIVER microscope.

## Data availability

Data underlying the results presented in this paper are not publicly available at this time but may be obtained from the authors upon request.

## Supplemental document

See Supplement 1 for supporting content.

## References

1. R. C. Arevalo, J. S. Urbach, and D. L. Blair, “Size-dependent rheology of type-I collagen networks,” Biophys. J. 99, 65–67 (2010).

2. R. C. Arevalo, P. Kumar, J. S. Urbach, and D. L. Blair, “Stress heterogeneities in sheared type-i collagen networks revealed by boundary stress microscopy,” PLoS One 10, 1–12 (2015).

3. M. J. Hogan, J. A. Alvarado, and J. E. Weddell, “Vitreous,” in Histology of the Human Eye: An Atlas and Textbook (W. B. Saunders Company, 1971), pp. 607–637.

4. S. L. Voytik-Harbin, A. O. Brightman, B. Z. Waisner, J. P. Robinson, and C. H. Lamar, “Small intestinal submucosa: A tissue-derived extracellular matrix that promotes tissue-specific growth and differentiation of cells in vitro,” Tissue Eng. 4, 157–174 (1998).

5. L. I. Los, M. J. A. Van Luyn, and P. Nieuwenhuis, “Organization of the rabbit vitreous body: Lamellae, Cloquet’s channel and a novel structure, the “alae canalis Cloqueti,”” Exp. Eye Res. 69, 343–350 (1999).

6. K. J. Bos, D. F. Holmes, R. S. Meadows, K. E. Kadler, D. McLeod, and P. N. Bishop, “Collagen fibril organisation in mammalian vitreous by freeze etch/rotary shadowing electron microscopy,” Micron 32, 301–306 (2001).

7. G. Eisner, “Zur Anatomie des Gaskorpers,” Albr. v. Graefes Arch. klim. exp. Ophthal. 193, 33–56 (1975).

8. J. A. Castrén, “Phase-contrast and electron microscopic studies of human and rabbit vitreous,” Acta Ophthalmol. 42, 651–656 (1964).

9. A. V. Noulas, A. D. Theocharis, E. Feretis, N. Papageorgakopoulou, N. K. Karamanos, and D. A. Theocharis, “Pig vitreous gel: Macromolecular composition with particular reference to hyaluronan-binding proteoglycans,” Biochimie 84, 295–302 (2002).

10. E. A. Balazs and J. L. Denlinger, “The Vitreus,” in The Eye, H. Davson, ed., 3rd ed. (Academic Press, 1984), pp. 533–589.

11. A. Buades, B. Coll, and J.-M. Morel, “Non-Local Means Denoising,” Image Process. Line 1, 208–212 (2011).

12. G. Steger, “An unbiased detector of curvilinear structures,” IEEE Trans. Pattern Anal. Mach. Intell. 20, 113–125 (1998).

13. C. J. Creveling, Y. Alsanea, and B. Coats, “Correlation of Collagen Fibril Properties and Inner Limiting Membrane Thickness with Vitreoretinal Adhesion in Human Eyes,” Exp. Eye Res. (2022).

14. P. Berens, “CircStat: A MATLAB Toolbox for Circular Statistics,” J. Stat. Softw. 31, 1–21 (2009).

15. J. Sebag and E. A. Balazs, “Morphology and ultrastructure of human vitreous fibers,” Investig. Ophthalmol. Vis. Sci. 30, 1867–1871 (1989).

16. S. P. Wright, “Adjusted P-Values for Simultaneous Inference,” Biometrics 48, 1005–1013 (1992).

17. E. A. Balazs, “Structure of the Vitreous Gel,” in Acta: XVII Concilium Ophthalmologicum 1954, International Congress of Ophthalmology, ed. (University of Toronto Press, 1955), pp. 1019–1024.

18. B. A. Filas, N. S. Shah, Q. Zhang, Y. B. Shui, S. P. Lake, and D. C. Beebe, “Quantitative imaging of enzymatic vitreolysis-induced fiber remodeling,” Investig. Ophthalmol. Vis. Sci. 55, 8626–8637 (2014).

19. C. S. Nickerson, J. Park, J. A. Kornfield, and H. Karageozian, “Rheological properties of the vitreous and the role of hyaluronic acid,” J. Biomech. 41, 1840–1846 (2008).

20. B. A. Filas, Q. Zhang, R. J. Okamoto, Y. B. Shui, and D. C. Beebe, “Enzymatic degradation identifies components responsible for the structural properties of the vitreous body,” Investig. Ophthalmol. Vis. Sci. 55, 55–63 (2014).

21. B. A. Bembridge, G. N. Crawford, and A. Pirie, “Phase-contrast microscopy of the animal vitreous body,” Br. J. Ophthalmol. 36, 131–142 (1952).

22. A. Dvornikov, L. Malacrida, and E. Gratton, “The DIVER microscope for imaging in scattering media,” Methods Protoc. 2, 1–12 (2019).

23. L. M. Jawerth, S. Munster, D. A. Vader, B. Fabry, and D. A. Weitz, “A Blind Spot in Confocal Reflection Microscopy□: The Dependence of Fiber Brightness on Fiber Orientation in Imaging Biopolymer Networks,” Biophys. J. 98, L01–L03 (2009).

24. J. Sebag, “Age-related changes in human vitreous structure,” Graefe’s Arch. Clin. Exp. Ophthalmol. 225, 89–93 (1987).

