## Supplement 1 for "Confocal reflectance microscopy for mapping collagen fiber organization in the vitreous gel of the eye"

© 2021 Optica Publishing Group under the terms of the [Optica Publishing Group Open Access Publishing Agreement](https://doi.org/10.1364/OA_License_v2)

*1.* *Illustration of decentered image acquisition*

To avoid the bright center spot artifact in the center of the optical imaging field and to minimize drift, we used the Olympus Fluoview software to limit image acquisition to a decentered subset (1/36th) of the optical imaging field.

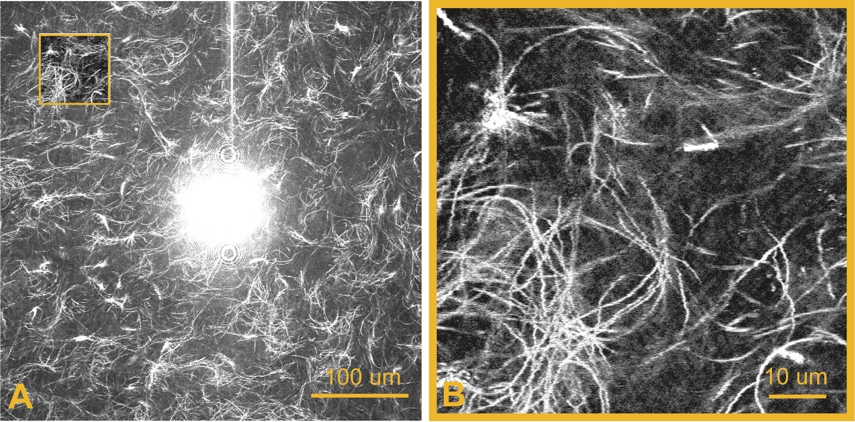

Fig. S1. Illustration of the bright spot artifact in the center of the optical imaging field (A), and image acquisition from a decentered part of the optical imaging field (orange box in A, B).

*2.* *Alternate image analysis methods for determining fiber density*

To assess the impact of the image analysis method on our findings, we applied two alternate image density analysis methods to the dataset from the anterior-posterior axis originating at the nasal limbus imaged in the tangential plane (Fig. S2). Both methods yielded statistically significant decreases in fiber density with increasing distance from inner cornea. Although automated fiber segmentation yielded higher values for fiber density than manual fiber counting (Fig. S2A, Fig 8A), correlation between the two methods was high (R = 0.90, p<0.001), supporting use both methods to assess relative differences in fiber density. Automated pixel thresholding also correlated well with automated fiber segmentation (R = 0.92, p<0.001).

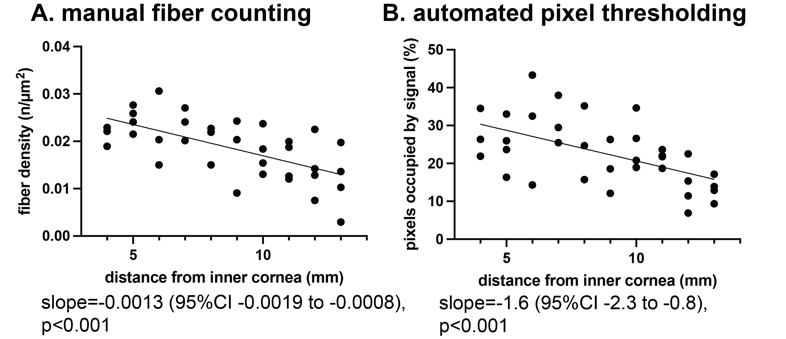

Fig. S2. Fiber density trend along the anterior-posterior axis originating at the nasal limbus using two methods of density quantification: A) manual fiber counting and B) automated pixel thresholding.

*3.* *Assessment of spatial variation of fiber width*

Fiber width was slightly, but significantly lower in the inferior dataset compared to the other four datasets (Fig. S3F). There was a small magnitude, statistically significant decrease in fiber width with increasing distance from inner cornea along the anterior-posterior axes originating at the superior and nasal limbus, but not along the axes originating at the inferior and temporal limbus (Fig. S3A-E).

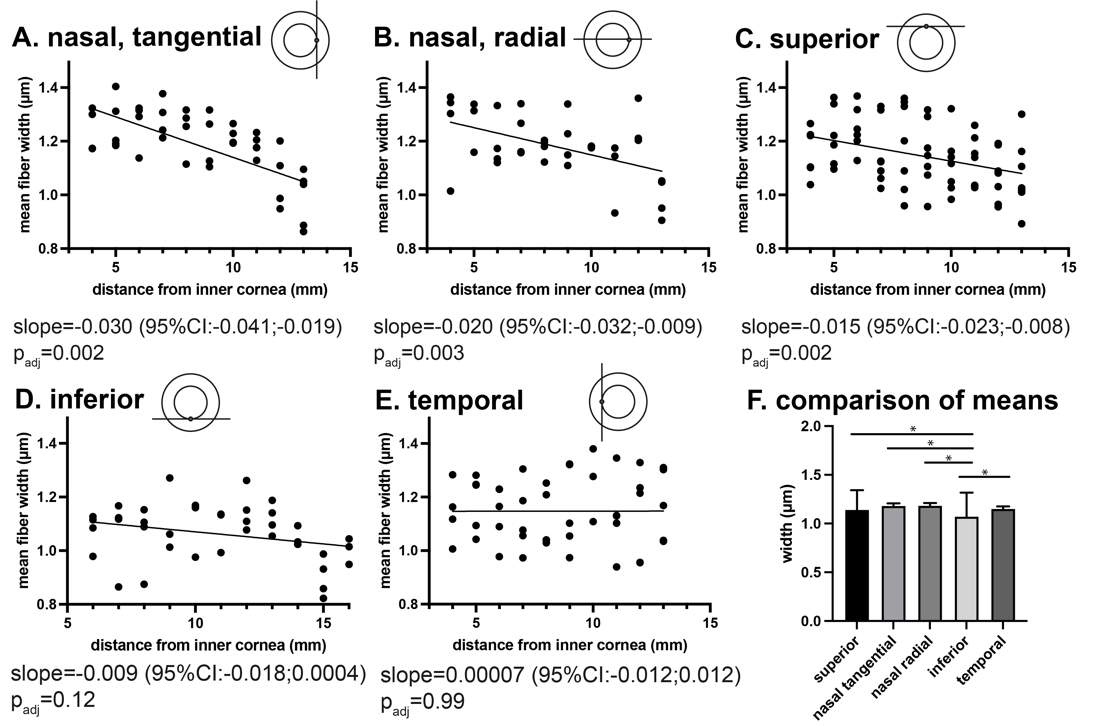

Fig S3. Trend in fiber width as measured by automated fiber segmentation along anterior-posterior axes originating at four different sites (A,B) nasal, (C) superior, (D) inferior, and (E) temporal and imaged in tangential planes (A, C, D, E) or a radial plane (B). F, comparison of overall width between the 5 datasets using a mixed effects model accounting for clustering and distance from inner cornea. Error bars represent standard error. Asterisks indicate p_adj_ ≤ 0.05.

Since the trends in width closely mirrored the trend in fiber density, we considered the possibility that there was overestimation of width at points of overlap. To evaluate this, we assessed correlation between mean width and fiber count in each individual image and found a strong correlation (R=0.87, Fig. S4).

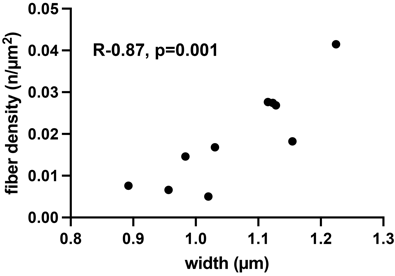

Fig S4. Correlation between mean fiber density and fiber count in each image in the dataset of images from the anterior-posterior axis originating at the nasal limbus imaged in the tangential plane.

*4.* *Assessment of spatial variation of in fiber alignment and orientation*

We evaluated whether there were any trends in alignment coefficient with increasing distance from inner cornea. We found a small increase in alignment coefficient from anterior to posterior along the superior limbus (Fig. S5C). When a radial plane was used to image the axis originating at the nasal limbus, we found a negative quadratic relationship (Fig. S5B), indicating significantly higher alignment coefficients in the center of the eye compared to the front and the back (p<0.001).

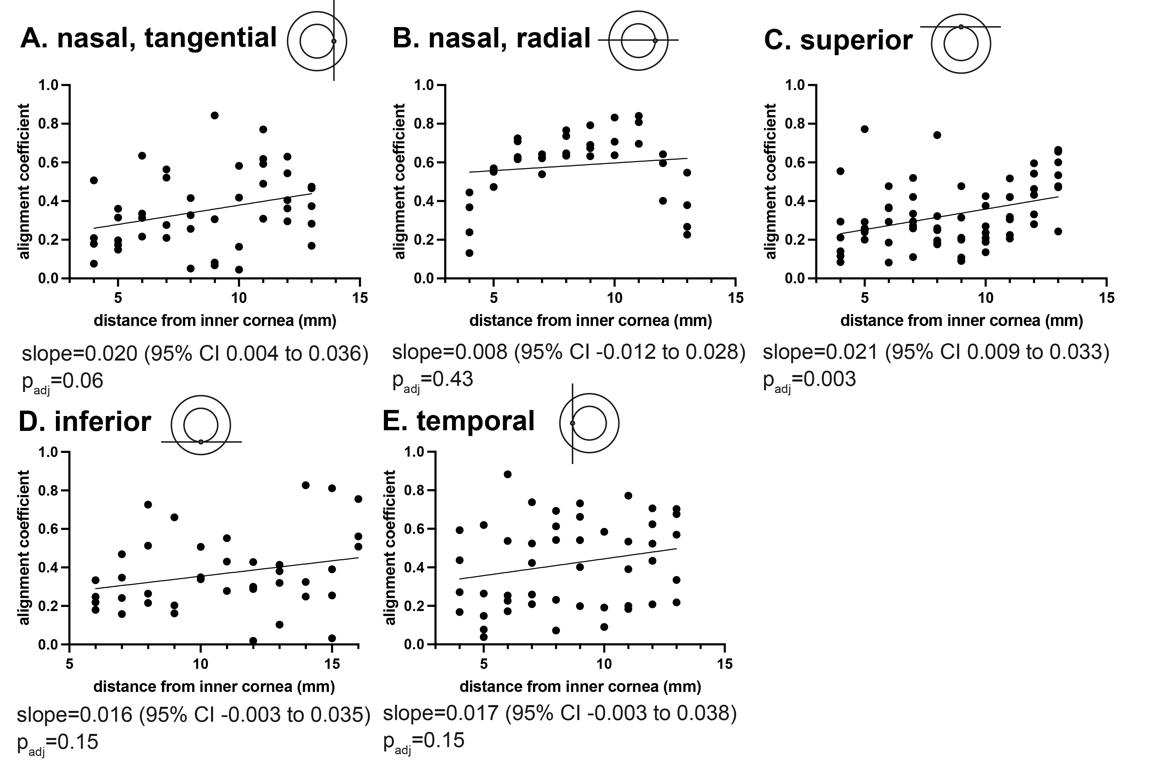

Fig. S5. Trend in alignment coefficient as measured by automated fiber segmentation along anterior-posterior axes originating at four different sites (A, B) nasal, (C) superior, (D) inferior, and (E) temporal and imaged in tangential planes (A, C, D, E) or a radial plane (B). Distance from the inner cornea is equivalent to distance along the A-P axis.

Visual inspection did not demonstrate any trends in angular mean with increasing distance from inner cornea along the anterior-posterior axes originating in any location (Fig. S6).
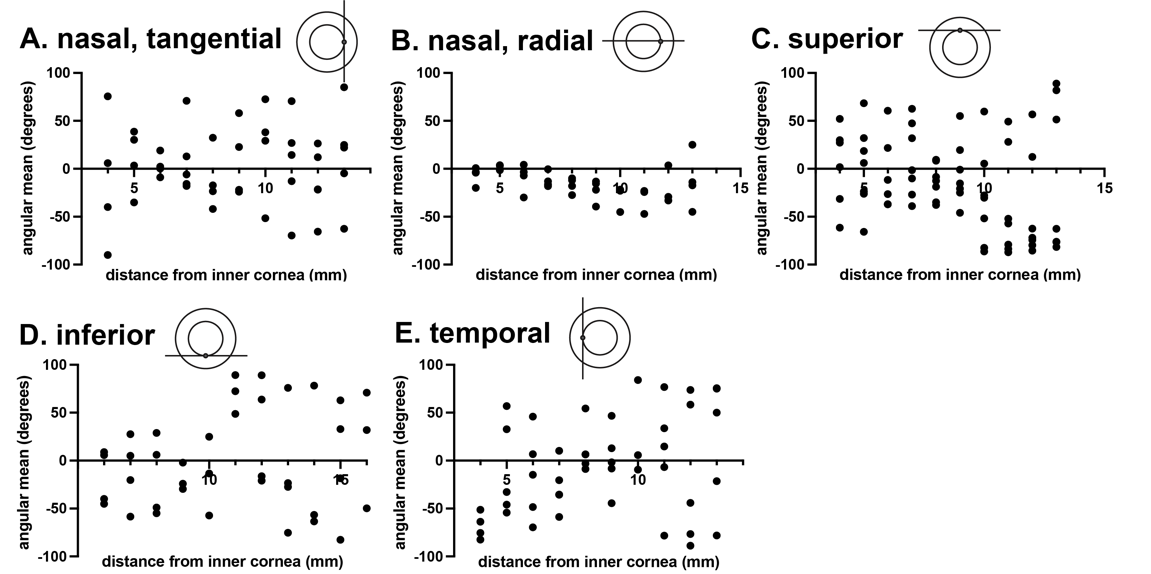

Fig S6. Trend in angular mean as measured by automated fiber segmentation along anterior-posterior axes originating at four different sites (A,B) nasal, (C) superior, (D) inferior, and (E) temporal and imaged in tangential planes (A, C, D, E) or a radial plane (B). Distance from the inner cornea is equivalent to distance along the A-P axis.

To verify the fiber alignment and orientation results from our automated fiber segmentation, we utilized a second method, OrientationJ, which relies upon first derivatives at each pixel, rather than fiber segmentation, to analyze data obtained with radial and tangential imaging planes along the anterior-posterior axis originating at the nasal limbus. OrientationJ outputs a dominant direction and coherency [17], which correspond to the angular mean and alignment coefficient, respectively. Similar to our prior findings, we found greater coherency when a radial plane was used to image the axis originating at the nasal limbus compared to when a tangential plane was used (Fig. S7). Also similar to our prior findings, the dominant direction in the radial plane clustered around 0, while the dominant direction in the tangential plane was widely distributed (Fig. S7).

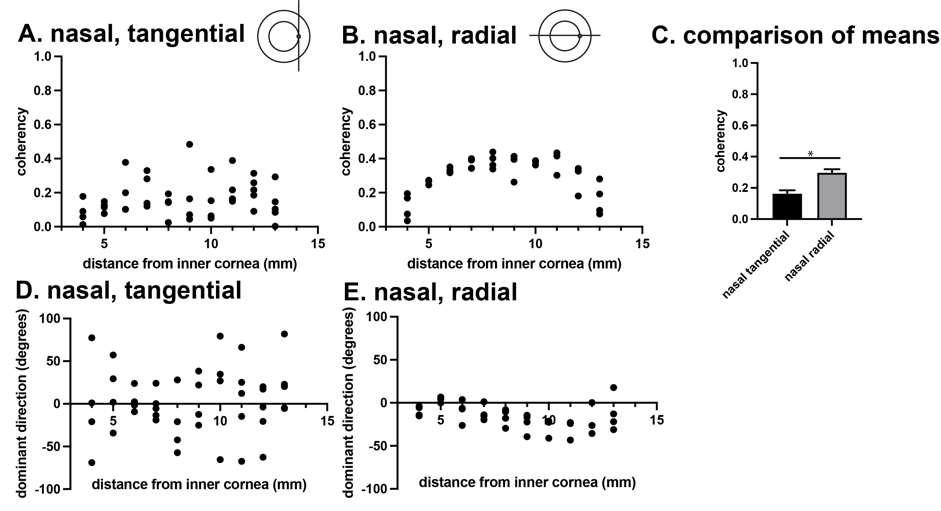

Fig S7. Trend in fiber coherency as measured by Orientation J along the anterior-posterior axis originating at the nasal limbus imagined in the tangential plane (A), and radial plane (B). Trend in dominant direction as measured by Orientation J along the anterior-posterior axis originating at the nasal limbus imagined in the tangential plane (C), and radial plane (D). F, comparison of coherency between the 2 datasets using a mixed effects model accounting for clustering and distance from inner cornea. Asterisk indicates p_adj_<0.001.

We evaluated correlation between the two methods of assessing the peak and spread of angles and found high correlation (R = 0.85, p=0.0017) between alignment coefficient and coherency, and high correlation (R = 0.98, p<0.0001) between angular mean and dominant direction.
